# Coronavirus S protein-induced fusion is blocked prior to hemifusion by Abl kinase inhibitors

**DOI:** 10.1101/246991

**Authors:** Jeanne M. Sisk, Matthew B. Frieman, Carolyn E. Machamer

## Abstract

Enveloped viruses gain entry into host cells by fusing with cellular membranes, a step required for virus replication. Coronaviruses, including the severe acute respiratory syndrome coronavirus (SARS-CoV), Middle East respiratory syndrome coronavirus (MERS-CoV), and infectious bronchitis virus (IBV), fuse at the plasma membrane or use receptor-mediated endocytosis and fuse with endosomes depending on the cell or tissue type. The virus Spike (S) protein mediates fusion with the host cell membrane. We have shown previously that an Abl kinase inhibitor, imatinib, significantly reduces SARS-CoV and MERS-CoV viral titers and prevents endosomal entry by HIV SARS S and MERS S pseudotyped virions. SARS-CoV and MERS-CoV are classified as BSL-3 viruses, which can make experimentation into the cellular mechanisms involved in infection more challenging. Here, we use IBV, a BSL-2 virus, as a model for studying the role of Abl kinase activity during coronavirus infection. We found that imatinib and two specific Abl kinase inhibitors, GNF2 and GNF5, reduce IBV titers by blocking the first round of virus infection. Additionally, all three drugs prevented IBV S-induced syncytia formation prior to the hemifusion step. Our results indicate that membrane fusion (both virus-cell and cell-cell) is blocked in the presence of Abl kinase inhibitors. Studying the effects of Abl kinase inhibitors on IBV will be useful in identifying host cell pathways required for coronavirus infection. This will provide insight into possible therapeutic targets to treat infections by current as well as newly emerging coronaviruses.

## INTRODUCTION

Coronaviruses are enveloped RNA viruses and include those that cause the common cold as well as the highly pathogenic SARS-CoV and MERS-CoV. In order to gain entry into host cells, all enveloped viruses, including coronaviruses, fuse with cellular membranes. Coronaviruses bind to cell surface receptors and fuse with the plasma membrane or are endocytosed and fuse with endosomes. This fusion step is required in order for the viral genome to be delivered to the host cell cytoplasm for replication. SARS and MERS coronaviruses fuse with late and early endosomes, respectively, and also at the plasma membrane (1–4). The virus S protein mediates fusion between viral and host cell membranes.

Since the outbreaks of SARS-CoV in 2003 and MERS-CoV in 2012, these viruses have been studied extensively in order to understand infection and pathogenicity with the goal of identifying therapeutic targets. A high-throughput screen identified an Abelson (Abl) kinase inhibitor, imatinib, as an inhibitor of SARS-CoV and MERS-CoV (5). Abl kinases are non-receptor tyrosine kinases localized to the cytoplasm. Two Abl kinases, Abl1 and Abl2, are expressed in human cells, and are involved in several cellular processes ranging from embryonic morphogenesis to mediating viral infections (6, 7). Imatinib is a small molecule inhibitor specifically designed to target the Abl kinase portion of the BCR-Abl protein (8), which results from a chromosomal translocation and causes chronic myeloid leukemia (CML) (9–11). Previously, we have shown that imatinib blocks entry of pseudotyped virions containing SARS-CoV or MERS-CoV S protein (12). Inhibition of SARS and MERS coronavirus entry by imatinib implicates Abl kinase activity in coronavirus infection, and reports from other groups demonstrate Abl kinase involvement in infections by several other viruses (13–20). This strongly suggests the Abl kinase signaling pathway is promising to investigate for development of antiviral therapies.

Based on results from our previous report, we hypothesize that Abl kinase activity is required for entry of other coronaviruses, presumably for virus-cell fusion. The classification of SARS-CoV and MERS-CoV as BSL-3 pathogens and SARS-CoV as a select agent, presents a barrier to the experimental methods that can be performed with the live viruses. Additionally, membrane composition of pseudotyped virions differs from that of true virions since pseudotyped virions bud from the plasma membrane while true virions bud from the endoplasmic reticulum-Golgi intermediate compartment (21, 22). This could lead to differences in virus-cell fusion and entry mechanisms. Because of this, it is important to evaluate these effects in a live virus system and it is advantageous to find a BSL-2 coronavirus that can be used as a model of infection. Here, we report the effects of imatinib and two other Abl kinase inhibitors, GNF2 and GNF5, on IBV infection. We show that the Abl kinase inhibitors interfere with the first round of IBV infection, indicating an inhibition of virus-cell fusion. In addition, we directly demonstrate that cell-cell fusion and syncytia formation mediated by IBV S is inhibited by imatinib, GNF2 and GNF5 prior to hemifusion, suggesting a role for Abl kinases in both virus-cell and cell-cell fusion events. IBV will be an excellent model to determine the mechanism of Abl kinase action during coronavirus entry in order to find new targets for therapeutic intervention.

## MATERIALS AND METHODS

### Cells and virus

Vero cells (ATCC CCL-81) were cultured in Dulbecco’s Modified Eagle Medium (DMEM; Invitrogen/Gibco, Grand Island, NY) supplemented with 10% fetal bovine serum (FBS; Atlanta Biologicals, Lawrenceville, GA) and 0.1 mg/ml Normocin (InvivoGen, San Diego, CA). Cells were maintained at 37°C with 5% CO_2_. The wild-type recombinant US Beaudette strain of IBV was used in all experiments (23).

### Plasmids

pCAGGS/SARS-CoV S and pCAGGS/VSV G were previously described (24, 25). Two codon optimized plasmids encoding portions of IBV Beaudette S1 and S2 were generously provided by Helene Verhije (Department of Pathobiology, Utrecht University). A full length, partially codon-optimized IBV S cDNA sequence was constructed in the mammalian expression vector pCAGSS as follows. Nucleotides 1-110 (numbering based on open reading frame) containing the signal sequence were PCR amplified with EcoRI and BstxI sites from a non-codon-optimized cDNA sequence previously cloned by reverse transcription-PCR using RNA from Vero cells infected with a Vero-adapted strain of IBV (26). This same plasmid was the template for PCR amplification of nucleotides 2956-3489 encoding the C-terminus flanked by XmaI and XhoI sites. Nucleotides 111-1613 flanked by BstxI and HindIII sites (and containing the RRFRR furin cleavage site between S1 and S2) were PCR amplified from a cDNA vector containing codon-optimized sequence of IBV S1, and nucleotides 1614-2955 flanked by HindIII and XmaI sites were PCR amplified from cDNA containing the codon-optimized sequence of IBV S2 (27, 28). QuikChange mutagenesis (Agilent Genomics) was used to remove the HindIII restriction site between S1 and S2 once the amplicons had been ligated together, and the full coding sequence was confirmed by dideoxy sequencing.

### Antibodies and Abl kinase inhibitors

The mouse monoclonal anti-IBV S antibody (9B1B6) recognizes the lumenal domain and was a kind gift from Ellen Collisson (Western University of Health Sciences) (29). The rabbit polyclonal anti-IBV S antibody to the C-terminus was previously described (26), as was the rabbit polyclonal antibody to SARS S (anti-S_CT_) (24). Rabbit anti-GFP (A-6455) was obtained from ThermoFisher. Alexa Fluor 488 anti-mouse IgG and Alexa Fluor 568 anti-rabbit IgG were obtained from Life Technologies (Grand Island, NY, USA). Cy5 anti-mouse IgG was obtained from Jackson Laboratories (West Grove, PA, USA). Imatinib (Cell Signaling Technologies, Danvers, MA), GNF2 and GNF5 (Selleckchem, Houston, TX) were diluted in DMSO and used at 10 µM in all experiments.

### IBV infection

Vero cells were seeded 5 × 10^5^ per dish in 35 mm dishes on coverslips 24 h prior to infection. Cells were pre-treated for 1 h prior to infection with 10 µM imatinib, GNF2 or GNF5, or an equal volume of DMSO, in serum- and antibiotic-free medium (SF). SF medium was removed and virus was added to cells in 200 µl total volume of SF medium −/+ drugs. Infections were carried out in medium containing 5% FBS −/+ drugs. After 8 or 18 h infections performed as described below, supernatants were harvested. Cellular debris was removed by a 3 min centrifugation at 2000 x g, 4°C, and directly used for plaque assays, or stored at −80°C. For infection at MOI 0.1 for 18 h, virus was adsorbed for 1 h at 37° C with gentle rocking every 10 min. Virus was removed and cells washed with 1 ml DMEM + 5% FBS. Fresh DMEM + 5% FBS was added to cells −/+ 10 µM imatinib, GNF2 or GNF5 and cells were incubated 18 h at 37°C. For infection at MOI 2 for 8h, cells were placed at 4° C for 10 min, and virus was adsorbed for 1 h at 4°C with gentle rocking every 10 min. Virus was removed and cells were washed with 1 ml DMEM + 5% FBS. Fresh DMEM + 5% FBS was added to cells −/+ 10 µM imatinib, GNF2 or GNF5 and cells were incubated 8 h at 37°C.

### Transient Transfection

The X-tremeGENE 9 DNA transfection reagent (Roche, Indianapolis, IN) was used to transiently transfect cells according to the manufacturer’s protocol. For all experiments, 35 mm dishes of Vero cells were transfected with 2 µg of pCAGGS/IBV S or pCAGGS/SARS-CoV S diluted into Opti-MEM medium (Invitrogen/Gibco, Grand Island, NY). Experiments were performed 24 h post-transfection unless otherwise indicated. For SARS CoV S, cells were treated for 15 min with 15 µg/ml trypsin at 37°C prior to examining syncytia formation as previously described (30).

### Indirect immunofluorescence microscopy

Cells were plated on coverslips in 35 mm dishes 24 h prior to infection or transfection, and infected or transfected as described above. To label total IBV S and SARS S proteins, at the indicated times post-infection or post-transfection, cells were fixed with 3% paraformaldehyde in PBS for 10 min at room temperature, permeabilized with block/perm buffer (PBS + 10% FBS, 0.05% Saponin, 10 mM glycine, 10 mM HEPES pH 7.4) and incubated with the appropriate primary antibodies (rabbit anti-IBV S diluted 1:500, rabbit anti-SARS S diluted 1:500 in block/perm buffer) for 15 min at room temperature. Cells were washed twice with PBS + 10 mM glycine and stained with secondary Alexa Fluor antibodies (diluted 1:1000 in block/perm buffer) for 15 min at room temperature. Cells were washed with PBS + glycine and DNA stained with Hoescht 33258. To label cell surface IBV S, intact cells were washed with ice-cold PBS, incubated 30 min on ice in block/perm buffer (without saponin) and incubated with mouse anti-IBV S (1:10) for 1 h on ice. Cells were then washed twice with ice-cold PBS, fixed and permeabilized as, and stained for total IBV S with rabbit anti-IBV S, and DNA as described above. Images were acquired with a Zeiss Axioscop microscope (Thornwood, NY) equipped for epifluorescence with a Sensys charge-coupled-device camera (Photometrics, Tucson, AZ), using IPLab software (Scanalytics, Vienna, VA). Images are shown inverted for better visualization of the IBV S, dsRed and YFP-GPI signals.

### Plaque assay

Vero cells were seeded at 5 × 10^5^ per dish in 35 mm dishes 24 h prior to infection. Supernatants from infected cells were serially diluted 10^−1^ through 10^−6^ in serum-free medium. Cells were washed with serum-free medium, 200 µl of diluted virus was added to each well and adsorption was allowed to proceed for 1 hour at 37°C with gentle rocking every 10 minutes. 2X DMEM and 1.6% agarose were mixed 1:1 and allowed to cool to 37°C. Cells were washed with SF medium, and 2 ml DMEM-agarose was added to each well, and cells were incubated for 72 hours at 37°C. DMEM-agarose was removed from each well, crystal violet staining was used to visualize plaques, and pfu/ml was calculated for each sample.

### Fusion Assay

HeLa and Vero cells were plated 3.5×10^5^ in 35 mm dishes 24 hr prior to transfection. HeLa cells were transfected as described above with 2 µg pCAGGS-IBV S and 0.5 µg of a plasmid encoding YFP-GPI anchor (Clontech) and Vero cells were transfected with 0.5 µg of a plasmid encoding dsRed (Clontech). Twenty-four h post-transfection, Vero cells were trypsinized and re-plated at 3.5 × 10^5^ in 35 mm dishes on coverslips and grown for 1 h in 10 µM imatinib, GNF2 or GNF5 and an equal volume of DMSO. An equal volume of HeLa cells was then added to each well of Vero and DMSO or drug was added to maintain treatment conditions. Twenty-four h later, cells were placed on ice. On ice, cells were incubated for 20 min in block/perm buffer as described previously (without saponin). A rabbit anti-GFP antibody and mouse anti-IBV S (1:500, 1 h on ice) were used to label surface YFP-GPI and IBV S, respectively. Cells were then fixed for 10 min in 3% PFA at RT, and all following steps were carried out at RT. Cells were washed with PBS and then permeabilized with block/perm buffer (0.05% saponin). Cells were washed and Alexa Fluor 488 anti-rabbit and Cy5 anti-mouse were used for secondary labeling of YPF-GPI and IBV S, respectively. Hoescht 33258 was used to stain DNA. All images were taken within 24 h with the same exposure as described above. Physical contact between Vero and HeLa cells was defined as an event. Each event was classified either as hemifusion or full fusion. For each treatment condition, total events were normalized to 100% and hemifusion events were normalized to the total number of events.

### IBV S processing analysis

Vero cells were plated 5×10^4^ per well in 35 mm dishes and 24 h later infected with IBV at MOI 0.1. Eighteen h post-infection supernatants were collected and virus was concentrated by layering 1 ml of supernatant onto 2 ml of a 20% sucrose cushion, and centrifuged for 1 h at 90K xg and 4°C in a TLA110 rotor. The supernatant was removed and samples resuspended and analyzed by SDS-PAGE on 4 to 12% acrylamide gradient gels (NUPAGE, ThermoFisher). Proteins were transferred to low fluorescence PVDF membrane (source) and blocked for 1 h in 5% non-fat dry milk in Tris-buffered saline with Tween (TBST) (150 mM NaCl, 10 mM Tris-Hcl [pH 7.4], 0.05% Tween-20). Membranes were incubated overnight at 4°C with rabbit anti-IBVS antibody 1:500 in 5% non-fat dry milk in TBST. Membranes were washed 3 × 5 min in TBST then incubated at room temperature for 1 h with goat anti-rabbit IgG 680 1:20,000 (Licor Biosciences, Lincoln, NE). Membranes were washed 3 × 5 min in TBST and rinsed in 1X TBS. Bands were visualized using the Licor Odyssey CLx and quantified using Image Studio software. The density of three IBV S bands, corresponding to S0, S2 and S2’, was measured and the percent for each band was calculated from the total S for each treatment condition.

## RESULTS

### IBV titers are reduced in infected Vero cells after treatment with Abl kinase inhibitors

We have shown previously that imatinib lowers SARS-CoV and MERS-CoV titers (12). We tested the effects of imatinib, as well as two other Abl kinase inhibitors, GNF2 and GNF5, on IBV infection in order to determine whether IBV could be used as a model to investigate Abl kinase involvement in coronavirus infection. We used two different types of inhibitors in order to determine target specificity as well mechanism of inhibition. Imatinib is a competitive inhibitor that binds near the ATP-binding pocket preventing binding of ATP (31, 32). Imatinib only binds when the Abl kinase is in the closed, inactive conformation (32). GNF2 and GNF5 bind to the auto-inhibitory myristate-binding site of the Abl kinase, leading to inactivation of the active kinase by promoting the closed conformation (33). Imatinib has several cellular targets, while GNF2 and GNF5 specifically inhibit the Abl kinases (33, 34). Treating infected cells with these specific inhibitors allowed us to evaluate the likelihood of Abl kinase involvement in coronavirus infection as opposed to any of the other cellular targets of imatinib. Vero cells were infected with IBV at MOI 0.1 in the absence and presence of the three inhibitors. Supernatants were harvested 18 h post-infection, and a plaque assay was used to measure viral titers of all samples. Results from this experiment showed that all three drugs significantly decreased virus titer. Imatinib lowered the viral titer ~90% when compared to control, and GNF2 and GNF5 lowered titer ~95% (Figure 1A).

**Figure 1.**
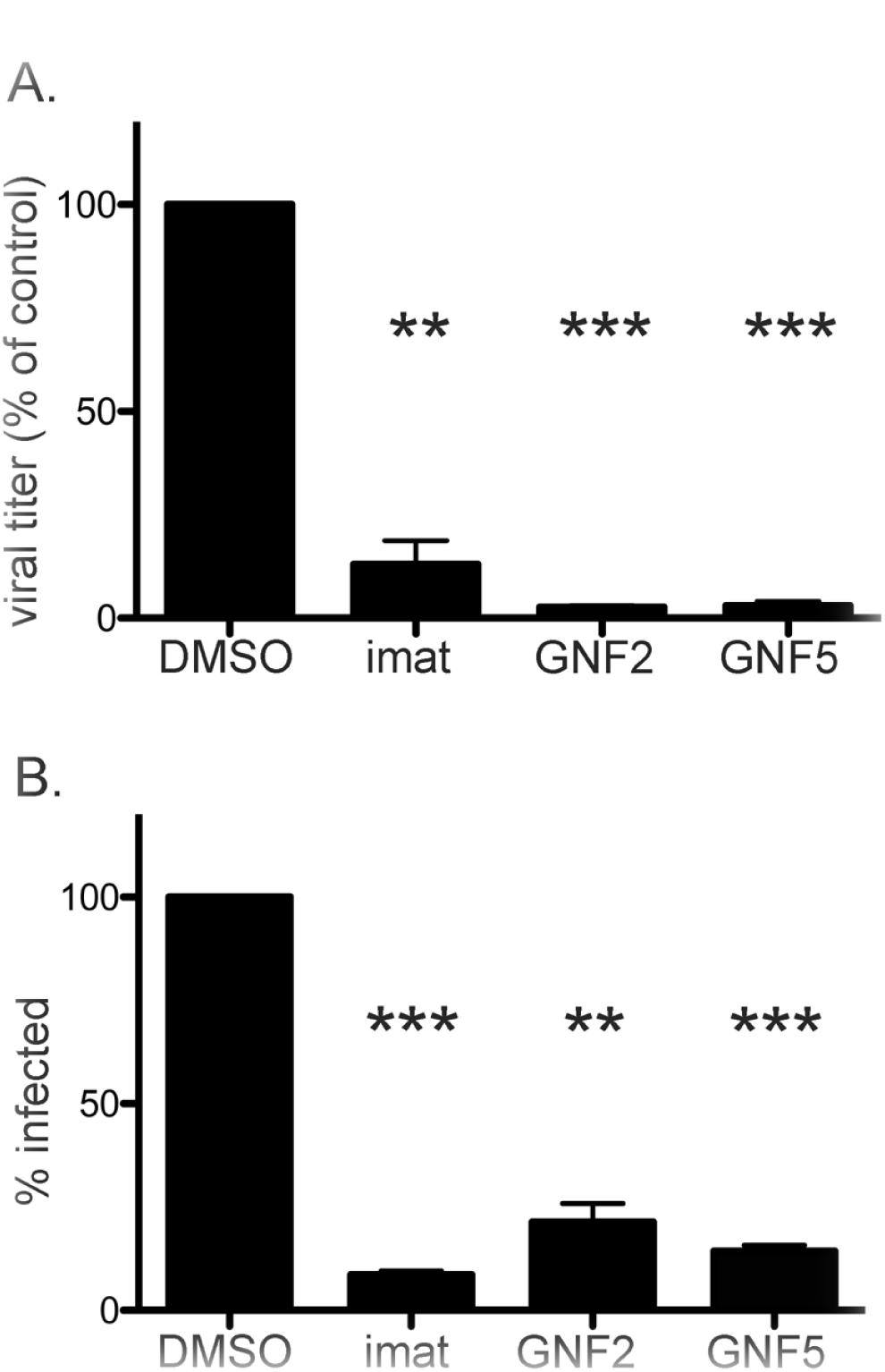
IBV infection is suppressed by Abl kinase inhibitors. Vero cells were pre-treated for 1 h with 10 µM imatinib, GNF2 or GNF5. Cells were adsorbed for 1 h with IBV at an MOI of 0.1 in the presence of DMSO or drug. (A) Supernatants were harvested and used for plaque assays to determine viral titer. (B) Immunofluorescence staining of the IBV S protein was used to count the number of infected cells in the presence of DMSO or drug. > 1,000 cells were analyzed per experimental condition in three independent experiments, and % of cells infected was calculated by comparing the number of infected cells to the number of total cells analyzed. Error bars represent standard error and p values were calculated using a paired t test. ** p < 0.01, *** p < 0.001.

Next, we determined whether the decrease in titer was due to reduced virus production in infected cells or the result of fewer cells being infected. Indirect immunofluorescence microscopy using an antibody to the IBV S protein was used to count the number of infected cells in the absence and presence of the Abl kinase inhibitors. Our observations showed that imatinib and GNF5 reduced the number of cells infected by ~90%, and GNF2 decreased the number of infected cells ~80% when compared to control (Figure 1B). These results demonstrate that the decrease in IBV titer was due to a decrease in the number of cells infected. The inhibition by GNF2 and GNF5 suggests Abl1 and/or Abl2 specifically play a role in IBV infection.

### Abl kinase inhibitors prevent the first round of IBV infection

Previously, we showed imatinib prevents entry of pseudotyped viruses expressing SARS-CoV or MERS-CoV S protein (12). In order to determine if IBV is blocked at the entry step by Abl kinase inhibitors, we infected Vero cells at a high MOI and for a duration that would only allow one round of infection. Supernatants were harvested at 8 h post-infection, and IBV titers were measured using a plaque assay. Results from this experiment showed a significant decrease in virus titer when IBV-infected cells were treated with imatinib, GNF2 or GNF5 as compared to control. Titer was reduced ~80% by imatinib and GNF2, and 70% by GNF5 (Figure 2A). We also observed a decrease in the total number of cells infected by 70% after treatment with all three drugs (Figure 2B).

**Figure 2.**
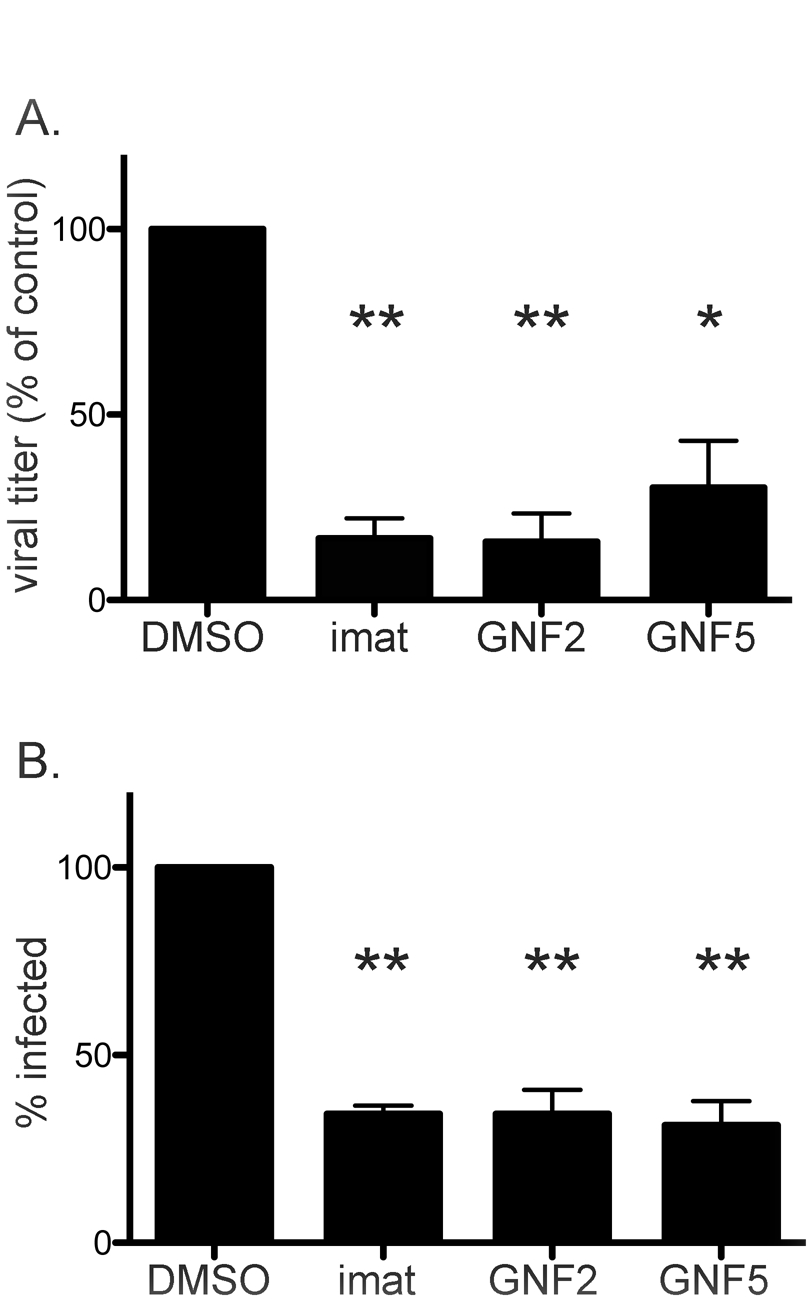
Abl kinase inhibitors block the first round of IBV infection. Vero cells were pre-treated for 1 h with DMSO, 10 µM imatinib, GNF2 or GNF5. Cells were adsorbed with IBV MOI 2 in the presence of DMSO or drug. Adsorption was performed at 4° C to allow synchronization of virus-receptor binding. Cells were washed and fresh medium containing DMSO or drug was added and infection was allowed to proceed for 8 h at 37° C. (A) Supernatants were harvested and used for plaque assays to determine viral titer. (B) Indirect immunofluorescence staining of the IBV S protein was used to count the number of infected cells in the presence of DMSO or drug. > 500 cells were analyzed per experimental condition in three independent experiments, and % of cells infected was calculated by comparing the number of infected cells to the number of total cells analyzed. Error bars represent standard error and p values were calculated using a paired t test. * p < 0.05, ** p < 0.01.

We considered that the decrease in number of cells infected could be due to decreased virus binding to host cell receptors. To address this, we infected Vero cells at a high MOI for 8 h and removed imatinib at indicated time points post-infection (Figure 3A). All cells were pre-treated with DMSO or imatinib, and either DMSO or imatinib was present in the infection medium during adsorption. Pre-treatment occurred from t-1 through t0, and adsorption occurred from t0 through t1. Imatinib is a reversible inhibitor of Abl kinases (35) and we observed a rapid reversal of its effects on IBV infection after its removal from the cell culture medium. Results from this experiment showed that when imatinib was removed from the cell culture medium immediately after adsorption, infection proceeded normally and viral titers matched those in untreated cells (Figure 3B t1, 3C imat 1h). However, when imatinib was present up to 1 h post-adsorption during virus fusion and entry, viral titers decreased significantly (Figure 3B t2, 3C imat 2 h). Similar to results in Figure 2, imatinib treatment for the duration of infection decreased IBV titers ~ 80% (Figure 3B). We also observed a decrease in the number of cells infected when imatinib was present in the cell culture medium for at least 1 h after adsorption (Figure 3C, imatinib 2 h). This result suggests that imatinib does not affect virus binding to host cell receptors, but does interfere with virus entry. The results from these experiments demonstrate that Abl kinase inhibitors prevent the first round of infection by IBV, potentially through interference with virus-cell fusion.

**Figure 3.**
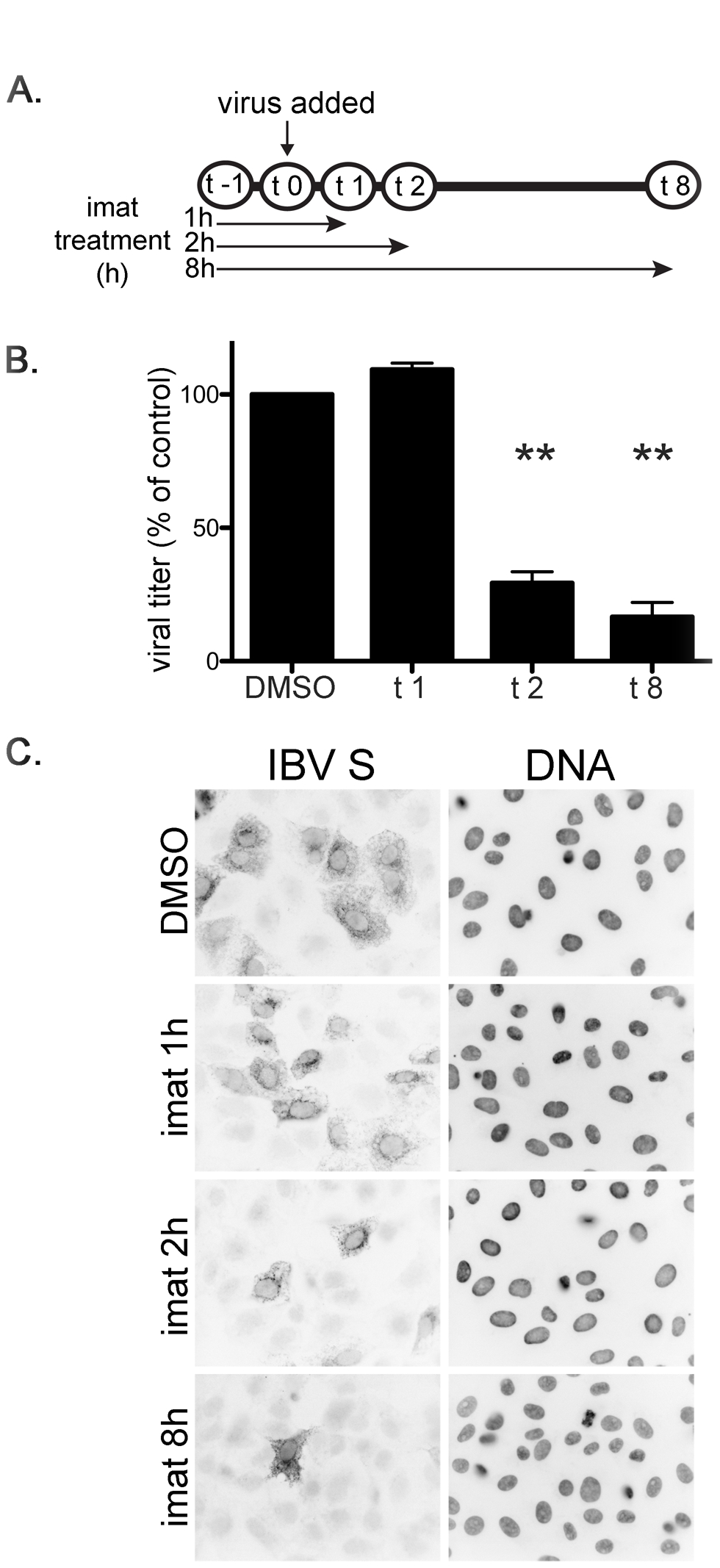
Imatinib interferes with the entry step of IBV infection. Vero cells were pre-treated for 1 h at 37° C with DMSO or 10 µM imatinib. Cells were adsorbed with IBV for 1 h at MOI 2 in the presence of DMSO or imatinib. Adsorption was performed at 4° C to allow synchronization of virus-receptor binding. Cells were washed and fresh medium containing DMSO or imatinib was added and infection was allowed to proceed for 8 h at 37° C. (A) Schematic showing duration of imatinib treatment. (B) Supernatants were harvested and used for plaque assays to determine viral titer. Results shown were calculated from three independent experiments. Error bars represent standard error and p values were calculated using a paired t test. ** p < 0.01. (C) Indirect immunofluorescence staining of the IBV S protein was used to visualize infected cells in the presence of DMSO or imatinib.

### Abl kinase inhibitors prevent virus-induced syncytia formation

Infection of Vero cells by IBV leads to cell-cell fusion and formation of multinucleated syncytia. Newly synthesized IBV S protein is incorporated into new virions that bud into the ER-Golgi intermediate compartment (22, 36), but the S protein is also transported to the cell surface late in infection. Just as S protein on the surface of the virion mediates virus-cell fusion, the S on the surface of infected cells mediates cell-cell fusion resulting in syncytia formation. Because the viral S protein mediates both of these events, it seems likely that virus-cell and cell-cell fusion may use the same cellular mechanism. Due to the difficulty in capturing individual virus-cell fusion events, we analyzed cell-cell fusion by measuring syncytium size in the absence and presence of imatinib, GNF2 or GNF5. In Vero cells that were infected with IBV, we observed syncytia formation with an average of 22 nuclei per syncytium (Figure 4A, B). Treatment with imatinib, GNF2 or GNF5 resulted in a statistically significant reduction of nuclei per syncytium. Average syncytium size was reduced to 5, 6 and 4 nuclei for imatinib, GNF2 and GNF5, respectively, an ~95% reduction in the presence of each drug (Figure 4A, B).

**Figure 4.**
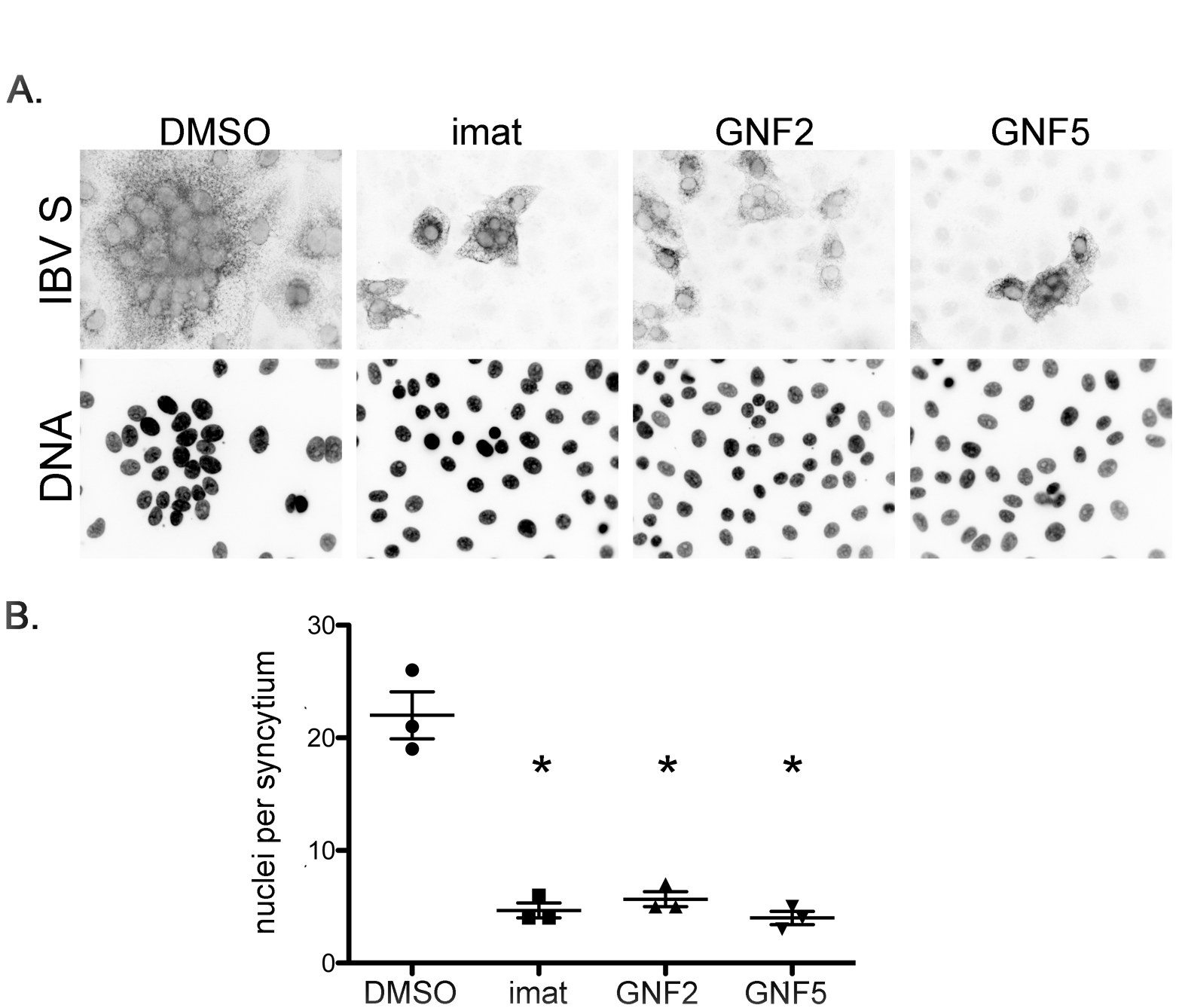
Abl kinase inhibitors prevent virus-induced syncytia formation in IBV-infected cells. Vero cells were pre-treated for 1 h at 37° C with DMSO, 10 µM imatinib, GNF2 or GNF5. Cells were adsorbed for 1 h with IBV MOI 0.1 in the presence of DMSO or drug. Cells were washed and fresh medium containing DMSO or drug was added and infection was allowed to proceed for 18 h at 37° C. (A) Indirect immunofluorescence assay detecting the S protein in infected cells. (B) Quantification of nuclei per syncytium in the presence or absence of Abl kinase inhibitors. ≥ 50 syncytia were analyzed in three independent experiments. Error bars represent standard deviation and p values were calculated using a paired t test. * p < 0.05.

Next, we determined whether imatinib, GNF2 and GNF5 inhibited syncytia formation induced by the S protein alone when exogenously expressed. Similar to infection, we observed formation of syncytia containing an average of 16 nuclei when Vero cells were transfected with a plasmid encoding IBV S (Figure 5A, B). Treating IBV S-transfected cells with Abl kinase inhibitors significantly reduced the number of nuclei per syncytium to ~4 nuclei for imatinib, GNF2 and GNF5, an ~80% reduction (Figure 5A, B). Because SARS-CoV is also inhibited by imatinib (12), we wanted to determine whether Abl kinase inhibitors interfere with fusion induced by SARS-CoV S. To this end, we also tested the effects of all three drugs on syncytia formation produced by the SARS-CoV S protein. Vero cells were transiently transfected with a plasmid encoding SARS-CoV S. The number of nuclei per syncytium was reduced 40% by imatinib, 50% by GNF2 or and 60% by GNF5, as compared to the number of nuclei per syncytium in untreated cells (Figure 5C, D). Average syncytium size induced by SARS-CoV S was much smaller than that induced by IBV S (7 nuclei and 16 nuclei per syncytium, respectively), but treatment with all three drugs did reduce the average to three nuclei per syncytium for both IBV S and SARS-CoV S. Note that treatment with Abl kinase inhibitors not only reduced the number of nuclei per syncytium, but also total number of syncytia. We did not quantify this observation, but while more than 50 syncytia containing greater than or equal to three nuclei were found in control cells, we routinely found fewer than 10 syncytia in drug-treated samples. Additionally, there are only two data points for the SARS-CoV S, imatinib-treated cells due to the fact that no syncytia were found in one of the replicate experiments. Treatment with imatinib, GNF2 or GNF5 had no effect on the total number of cells expressing VSV-G after transfection of a control plasmid (pCAGGS/VSV-G), demonstrating that the effects of these inhibitors were not due to transfection efficiency (data not shown). Together, these results demonstrate that imatinib, GNF2 and GNF5 interfere with syncytia formation mediated by the S protein both during IBV infection and when S protein is exogenously expressed in the absence of infection. The fact that we see the same phenotype with SARS-CoV S and IBV S reinforces the idea that IBV can be used as a model coronavirus to study Abl kinase involvement in coronavirus infection.

**Figure 5.**
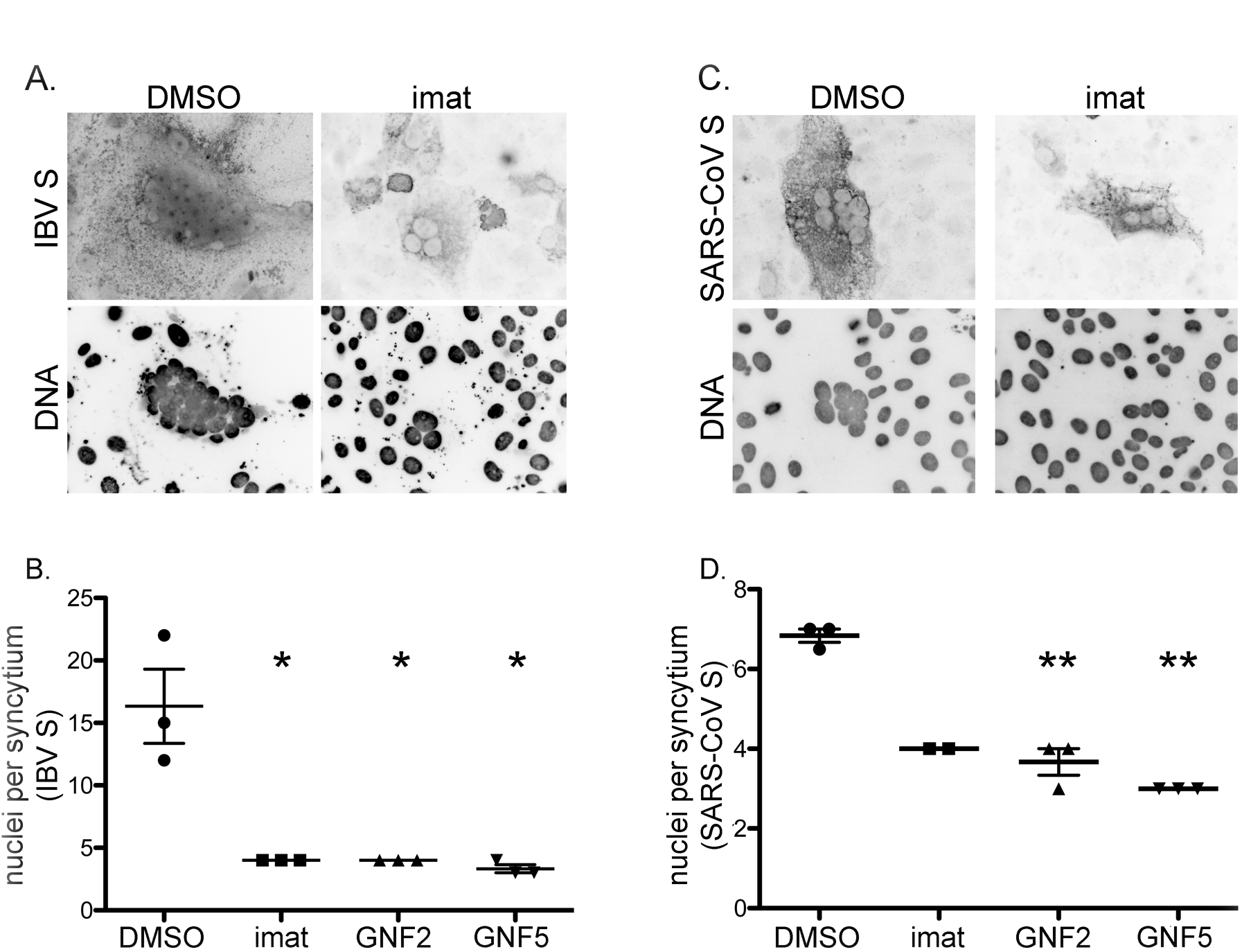
Abl kinase inhibitors prevent syncytia formation in cells exogenously expressing coronavirus S proteins. Vero cells were transfected with a plasmid encoding IBV S (A,B) or SARS-CoV S (C,D) and DMSO or 10 µM imatinib, GNF2 or GNF5 was added to the medium 3 h post-transfection. Cells were examined 48 h post-transfection. Top images show indirect immunofluorescence assay detecting IBV S (A) or SARS-CoV S (C). Graphs show quantification of nuclei per syncytium for IBV S (B) or SARS-CoV S (D) in the presence of DMSO or drug. (B, D) > 50 syncytia were analyzed in each of three independent experiments. Error bars represent standard deviation and p values were calculated using a paired t test. * p < 0.05, ** p < 0.01.

### Imatinib, GNF2 and GNF5 block IBV S-induced fusion prior to hemifusion

Virus fusion with host cells and virus-induced cell-cell fusion occurs in two steps. Mixing of the outer membrane leaflets (hemifusion) is followed by inner leaflet mixing and complete fusion. A previous report demonstrated that imatinib inhibits HIV env-induced cell-cell fusion at a post-hemifusion step, as indicated by an accumulation of hemifusion events observed in imatinib-treated cells (15). To determine whether imatinib, GNF2 and GNF5 inhibit IBV S-induced cell-cell fusion pre-or post-hemifusion, we used an assay similar to that used for HIV to distinguish hemifusion from full fusion events. HeLa cells were transfected with one plasmid encoding IBV S and a second encoding YFP targeted to the outer leaflet of the plasma membrane by a glycosylphosphatidylinositol anchor (YFP-GPI). After transfection, these cells were co-cultured with Vero cells transfected with a plasmid expressing soluble dsRed. Hemifusion between a HeLa and a Vero cell results in both cells containing the GPI anchor at the plasma membrane, with cytoplasmic dsRed signal observed in only one of the two cells. Full cell-cell fusion results in both cells with cytoplasmic dsRed signal and containing the labeled GPI anchor at the plasma membrane. Fusion events were observed and categorized as full fusion or hemifusion (Figure 6A). IBV S staining is not shown, but was used to ensure coexpression of IBV S and YFP-GPI in HeLa cells. We observed 54 total fusion events in the control sample and 25, 20 and 24 total fusion events in samples treated wtih imatinib, GNF2 and GNF5 samples, respectively. Hemifusion events accounted for 14% of all fusion events from control cells, and 8%, 11% and 12%, in cells treated with imatinib, GNF2 or GNF5, respectively (Figure 6B). Although total fusion events in drug-treated cells were reduced as expected, treatment with imatinib, GNF2 or GNF5 showed no significant change in the percent of hemifusion events as compared to control cells (Figure 6B), suggesting that Abl kinase inhibitors block the initial hemifusion step. This result demonstrates that Abl kinase inhibitors interfere with the cell-cell fusion necessary for syncytia formation, and provides evidence that these drugs may inhibit virus-cell fusion by preventing hemifusion, the first step in membrane mixing.

**Figure 6.**
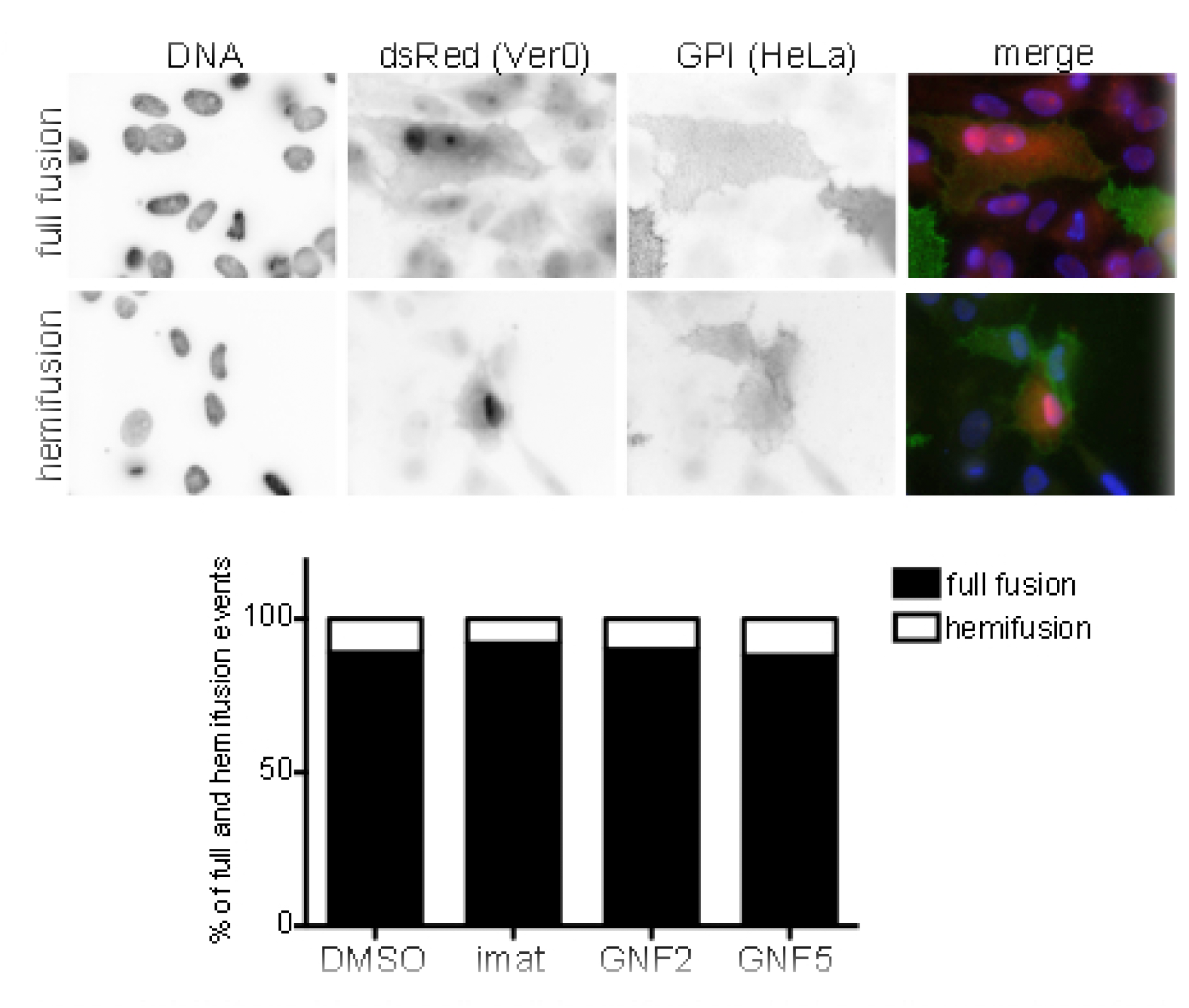
Abl kinase inhibitors block cell-cell hemifusion. HeLa cells were transfected with plasmids encoding IBV S and YFP-GPI. Vero cells were transfected with a plasmid encoding dsRed. Cells were co-cultured 24 h post-transfection, −/+ drug. IBV S, YFP-GPI and dsRed were detected 24 h later. Surface IBV S (not shown) was labeled with mouse anti-IBV S and Cy5 anti-mouse IgG. Surface YFP-GPI was labeled with rabbit anti-GFP and Alexa 488 anti-rabbit IgG. (A) Fusion events were analyzed and categorized as fusion (top panels) or hemifusion (bottom panels). (B) Quantification of hemifusion and full fusion events from two independent experiments. Total fusion events in control and drug-treated cells were normalized to 100% and the percent of hemifusion events was calculated based on comparison to total events for each treatment condition.

### Abl kinase inhibitors do not affect IBV S processing

We considered that treatment with Abl kinase inhibitors could affect IBV S processing and that this could be the cause of the block in cell-cell fusion. The full length S protein (S0) undergoes two cleavage events necessary for cell-cell fusion to take place. The first occurs at the S1/S2 site and produces two fragments, S1 and S2 (2). The second cleavage occurs at the S2’ site, resulting in exposure of the fusion peptide (3, 4). In the strain of Vero cells used here, IBV S is cleaved during virion exocytosis, providing a fusion-competent virus prior to host cell receptor binding. Because exposure of the fusion peptide has been shown to be necessary for cell-cell fusion and syncytia formation, we measured the amounts of each cleavage product after treatment with imatinib, GNF2 or GNF5. None of the drug treatments decreased the relative amounts of S0, S2 or S2’ (Figure 7A, 7B), indicating that IBV S protein processing and cleavage are not affected by Abl kinase inhibitors.

**Figure 7.**
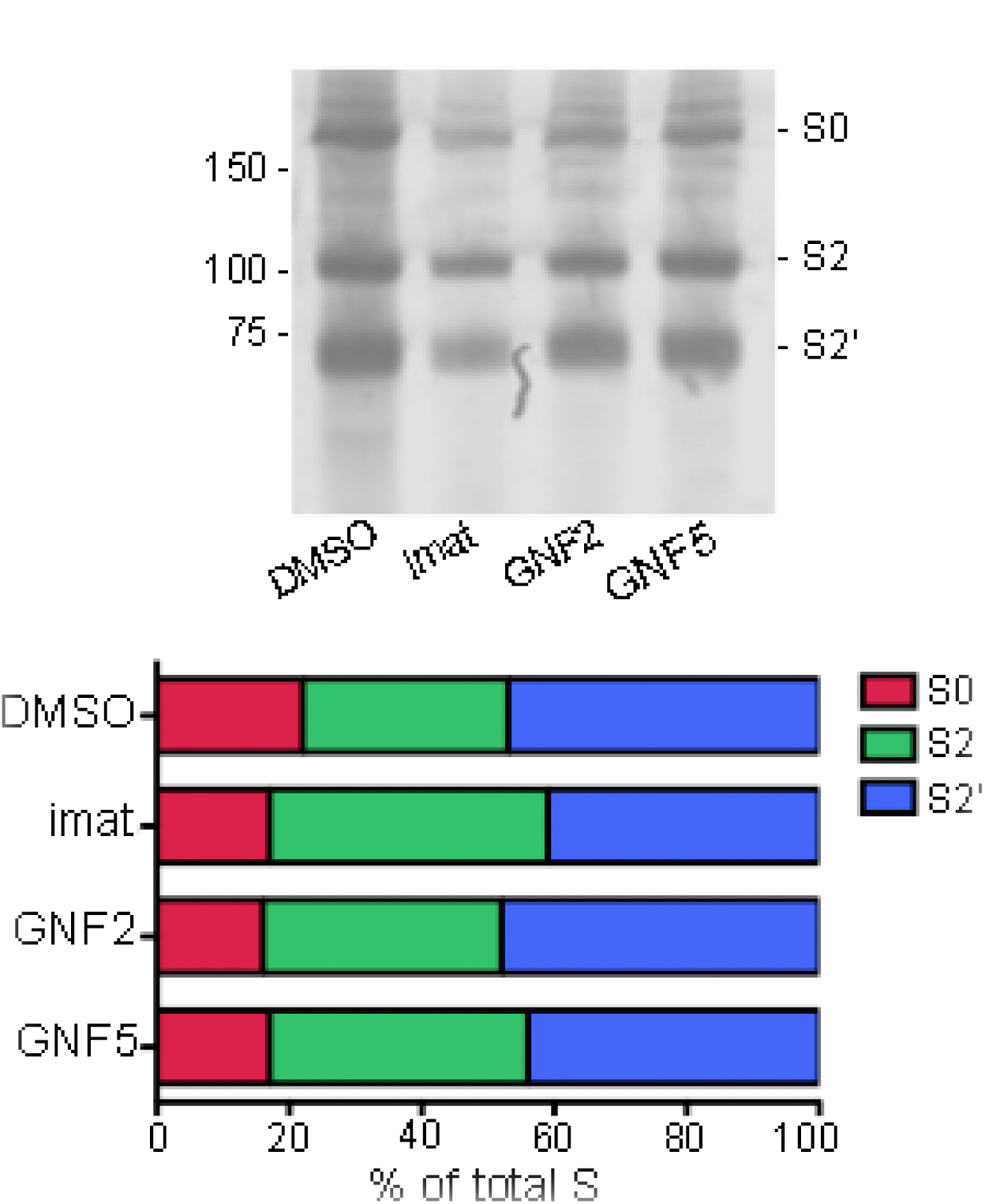
Abl kinase inhibitors do not affect IBV S processing. Vero cells were pretreated for 1 h with DMSO or 10 µM imatinib, GNF2 or GNF5, then infected with IBV at MOI 0.1 for 18 h, −/+ DMSO or 10 µM imatinib, GNF2 or GNF5. (A) A sucrose cushion was used to concentrate virus from supernatants of control and drug-treated IBV-infected cells. Concentrated virus was analyzed by SDS-PAGE and western blotting using rabbit anti-IBV S (1:500) and Licor anti-rabbit IgG 680 (1:10,000). (B) The amount of S0, S2 and S2’ was calculated by normalizing signal to the total amount of S within each sample. Results were confirmed in two independent experiments.

## DISCUSSION

We have demonstrated here that the Abl kinase inhibitors, imatinib, GNF2 and GNF5 reduce the viral titer of IBV by decreasing the number of cells infected, and that the first round of virus infection is inhibited. Our data show that each of the three inhibitors prevents syncytia formation induced by IBV as indicated by fewer nuclei per syncytium as compared to infected, untreated cells. A decrease in nuclei per syncytium in IBV S- and SARS-CoV S-expressing cells treated with imatinib, GNF2 or GNF5 demonstrates Abl kinase inhibitors prevent syncytia formation induced by the S protein in the absence of other viral proteins. Our results demonstrate that fusion was inhibited before the hemifusion step, prior to any lipid mixing of membranes. Additionally, we show that cleavage of S into S2 and S2’ was not affected by imatinib, GNF2 or GNF5, suggesting that IBV S processing and cleavage is independent of Abl kinase activity. The inhibition by specific inhibitors of Abl kinases, GNF2 and GNF5, strongly suggests that IBV infection requires the activity of Abl1 and/or Abl2. Our previous report showed that Abl2, but not Abl1, is required for entry of SARS-CoV and MERS-CoV pseudovirions (12), and we hypothesized that the inhibition of SARS-CoV and MERS-CoV infection was due to a block in virus fusion with the endosome membrane. Here we show definitively that Abl kinase inhibitors block coronavirus S-mediated fusion.

Abl1 and Abl2 are part of a kinase superfamily including Src kinase. These proteins contain conserved N-terminal domains, Src-homology domain 2 (SH2), Src-homology domain 3 (SH3) and tyrosine kinase domains. Additionally, they contain an F-actin-binding domain (FABD) connected by a linker region. Abl1 contains one FABD, while Abl2 contains two. Abl kinases are involved in numerous cellular functions providing communication between cell signaling and actin cytoskeleton organization (37). They are involved in T cell signaling (38, 39), embryonic development and (6, 40), and cell migration and invasion in cancer (41, 42). The distinct domains provide Abl kinases with the ability to act as scaffolds to facilitate building of signaling complexes (6, 43, 44), and to regulate protein function through phosphorylation of downstream protein targets (43). Binding actin through the FABD directly regulates cytoskeleton dynamics (44–47) to activate the essential cellular pathways described above. We hypothesize that coronavirus infection requires the multi-functional actions of Abl kinases for virus fusion at the plasma membrane or endosome membrane in order to gain entry into host cells.

It is likely that virus-cell and cell-cell fusion induced by the coronavirus S protein use the same or a very similar mechanism. Because Abl kinase activity plays a role in cytoskeletal rearrangement (47, 48), we hypothesize that Abl kinase inhibitors act by interfering with actin dynamics required for virus-cell and cell-cell fusion. Our hypothesis is supported on two fronts. First, cytoskeletal dynamics have been implicated in infection by several viruses (13–20). Actin cytoskeleton rearrangement was shown to be required for infection by Coxsackievirus (13), Vaccinia virus (14), HIV (15, 16), and Variola and monkeypox (17). These studies demonstrate that cytoskeletal rearrangement is a common mechanism required by many cellular processes that can be hijacked by viruses during infection, as reviewed by others (7, 49–51). Second, Abl kinase’s role in actin cytoskeletal rearrangement is required for essential cell-cell fusion events, including but not limited to, osteoclast (52) and myoblast fusion (53, 54) during development. Actin-induced podosome like structures are required for fusion of osteoclasts and myoblasts, presumably to alter membrane tension (55, 56). Interestingly, Harmon et al. (15) reported that cell-cell fusion mediated by the HIV envelope protein was inhibited by imatinib at a post-hemifusion step. We did not observe an increase in hemifusion events after treatment of IBV-infected cells with imatinib, GNF2 or GNF5. Since we used a different virus, cell line and fusion assay, it is difficult to compare the two results. We favor the idea that virus interaction with host cell receptors induces actin rearrangements through Abl kinase activity, and that this brings cell membranes in close proximity to allow hemifusion followed by full fusion (55).

Our results demonstrate that the coronavirus-membrane fusion step is blocked by Abl kinase inhibitors, as suggested by our previous report (12). Our results also indicate that Abl kinase activity may be involved in the membrane fusion events required for infection by all coronaviruses, and that imatinib, GNF2 and GNF5 have the potential to be general coronavirus inhibitors. This work strongly suggests that proteins involved in the Abl kinase pathway are excellent targets for developing treatments for coronavirus infections. Current studies are underway to determine the Abl kinase functions involved in coronavirus infection as well as determining whether the activity of Abl1 or Abl2, or both kinases is required for coronavirus infection. Given the role Abl kinases play in infection by so many other viruses makes this pathway a prime target for use in developing broadly acting antiviral therapeutics.

## ACKNOWLEDGEMENTS

We thank Helene Verhije (Department of Pathobiology, Utrecht University) for the kind gift of plasmids used to construct the codon-optimized IBV S, Jason Westerbeck for assembling the full-length IBV S clone, and Ellen Collison (Western University of the Health Sciences) for monoclonal anti-IBV S antibodies. We also thank Catherine Gilbert, Jason Westerbeck and Thiagarajan Venkataraman for helpful comments on the manuscript.

This was supported by HHS | NIH | National Institute of Allergy and Infectious Diseases (NIAID) AI118303 (JMS) and R01AI095569 (MBF).

